## Supplement for "Predicting chromatin conformation contact maps"

| Rank | Loss | Assay | Cell type | Position | Layers | Nodes | Dropout | Learning Rate |
|---|---|---|---|---|---|---|---|---|
| 1 | 0.0523 | 128 | 16 | 128 | 4 | 256 | 0.4 | 0.0005 |
| 2 | 0.0524 | 64 | 64 | 512 | 6 | 1024 | 0.2 | 0.0050 |
| 3 | 0.0527 | 64 | 128 | 128 | 4 | 1024 | 0.2 | 0.0050 |
| 4 | 0.0529 | 128 | 16 | 1024 | 4 | 256 | 0.2 | 0.0005 |
| 5 | 0.0531 | 256 | 256 | 128 | 6 | 1024 | 0.4 | 0.0005 |
| 6 | 0.0534 | 16 | 256 | 1024 | 2 | 1024 | 0.2 | 0.0005 |
| 7 | 0.0538 | 128 | 16 | 128 | 8 | 1024 | 0.4 | 0.0005 |
| 8 | 0.0539 | 256 | 128 | 128 | 4 | 256 | 0.2 | 0.0050 |
| 9 | 0.0540 | 128 | 64 | 256 | 4 | 256 | 0.6 | 0.0005 |

Table S1: **Results of the hyperparameter search.** The table lists the best-performing nine hyperparameter settings, with the corresponding loss values.

| Biosource | Assay Type | Accession | Generating Lab |
|---|---|---|---|
| 192627 | Dilution Hi-C | 4DNESYPKLMAM | Erez Lieberman Aiden |
| 192627 | in situ Hi-C | 4DNESECNR4O8 | Erez Lieberman Aiden |
| CC-2551 | Dilution Hi-C | 4DNESUB35TII | Erez Lieberman Aiden |
| CC-2551 | in situ Hi-C | 4DNESIE5R9HS | Erez Lieberman Aiden |
| GM12878 | Dilution Hi-C | 4DNESLLTENG9 | Bing Ren |
| GM12878 | DNA SPRITE | 4DNESI1U7ZW9 | Mitchell Guttman |
| GM12878 | in situ ChIA-PET CTCF protein | 4DNES7IB5LY9 | Yijun Ruan |
| GM12878 | in situ ChIA-PET RNA Pol II | 4DNESZ25MOZV | Yijun Ruan |
| GM12878 | in situ Hi-C | 4DNESPXW8XHY | Erez Lieberman Aiden |
| GM12878 | PLAC-seq H3K4me3 | 4DNESL3LFLGI | Bing Ren |
| H1-hESC | in situ ChIA-PET CTCF protein | 4DNESR9S8R38 | Yijun Ruan |
| H1-hESC | in situ ChIA-PET RNA Pol II | 4DNESNYUGLUN | Yijun Ruan |
| H1-hESC | in situ Hi-C | 4DNES2M5JIGV | Job Dekker |
| H1-hESC | Micro-C | 4DNES21D8SP8 | Job Dekker |
| H1-hESC | PLAC-seq H3K4me3 | 4DNESQMO66LZ | Bing Ren |
| HeLa cell line | Dilution Hi-C | 4DNESWMJBQMR | Jan-Michael Peters |
| HeLa cell line | DNase Hi-C | 4DNESGEEV6TJ | Todd Waldman |
| HeLa cell line | in situ Hi-C | 4DNESEDV9YMX | Job Dekker |
| HeLa cell line | Micro-C | 4DNESA5PN8AB | Job Dekker |
| HFF-hTERT | Dilution Hi-C | 4DNES9L4AK6Q | Job Dekker |
| HFF-hTERT | in situ Hi-C | 4DNESB6MNCFE | Job Dekker |
| HFF-hTERT | Micro-C | 4DNESEGKQY7I | Job Dekker |
| HFFc6 (Tier 1) | DNA SPRITE | 4DNESJYGTI8S | Mitchell Guttman |
| HFFc6 (Tier 1) | in situ ChIA-PET CTCF protein | 4DNESCQ7ZD21 | Yijun Ruan |
| HFFc6 (Tier 1) | in situ ChIA-PET RNA Pol II | 4DNESI1WZ5HT | Yijun Ruan |
| HFFc6 (Tier 1) | in situ Hi-C | 4DNES2R6PUEK | Job Dekker |
| HFFc6 (Tier 1) | Micro-C | 4DNESWST3UBH | Job Dekker |
| HFFc6 (Tier 1) | PLAC-seq H3K4me3 | 4DNESIF5UIQE | Bing Ren |
| HUVEC cell | Dilution Hi-C | 4DNESOSE2FYZ | Erez Lieberman Aiden |
| HUVEC cell | in situ Hi-C | 4DNESEW5JLUC | Erez Lieberman Aiden |
| IMR-90 | Dilution Hi-C | 4DNESSM1H92K | Erez Lieberman Aiden |
| IMR-90 | in situ Hi-C | 4DNES1ZEJNRU | Erez Lieberman Aiden |
| WTC-11 | in situ ChIA-PET CTCF protein | 4DNES8MZ76GP | Yijun Ruan |
| WTC-11 | in situ ChIA-PET RNA Pol II | 4DNESRRTL4BU | Yijun Ruan |
| WTC-11 | in situ Hi-C | 4DNESPDEZNWX | Job Dekker |
| WTC-11 | Micro-C | 4DNESODGV2V2 | Job Dekker |
| WTC-11 | PLAC-seq H3K4me3 | 4DNESDRL4ZKM | Bing Ren |
| WTC-11 AAVS1-GFP C28 | DNase Hi-C | 4DNES8BLXVP5 | Chuck Murry |
| WTC-11 AAVS1-GFP C28 | in situ Hi-C | 4DNESJ7S5NDJ | Job Dekker |
| WTC-11 AAVS1-GFP C28 | Micro-C | 4DNESAGG7EUC | Job Dekker |
| WTC-11 AAVS1-GFP C28 | PLAC-seq H3K4me3 | 4DNESIZ5TTHO | Bing Ren |

Table S2: **4D Nucleome data sets used in this study.**

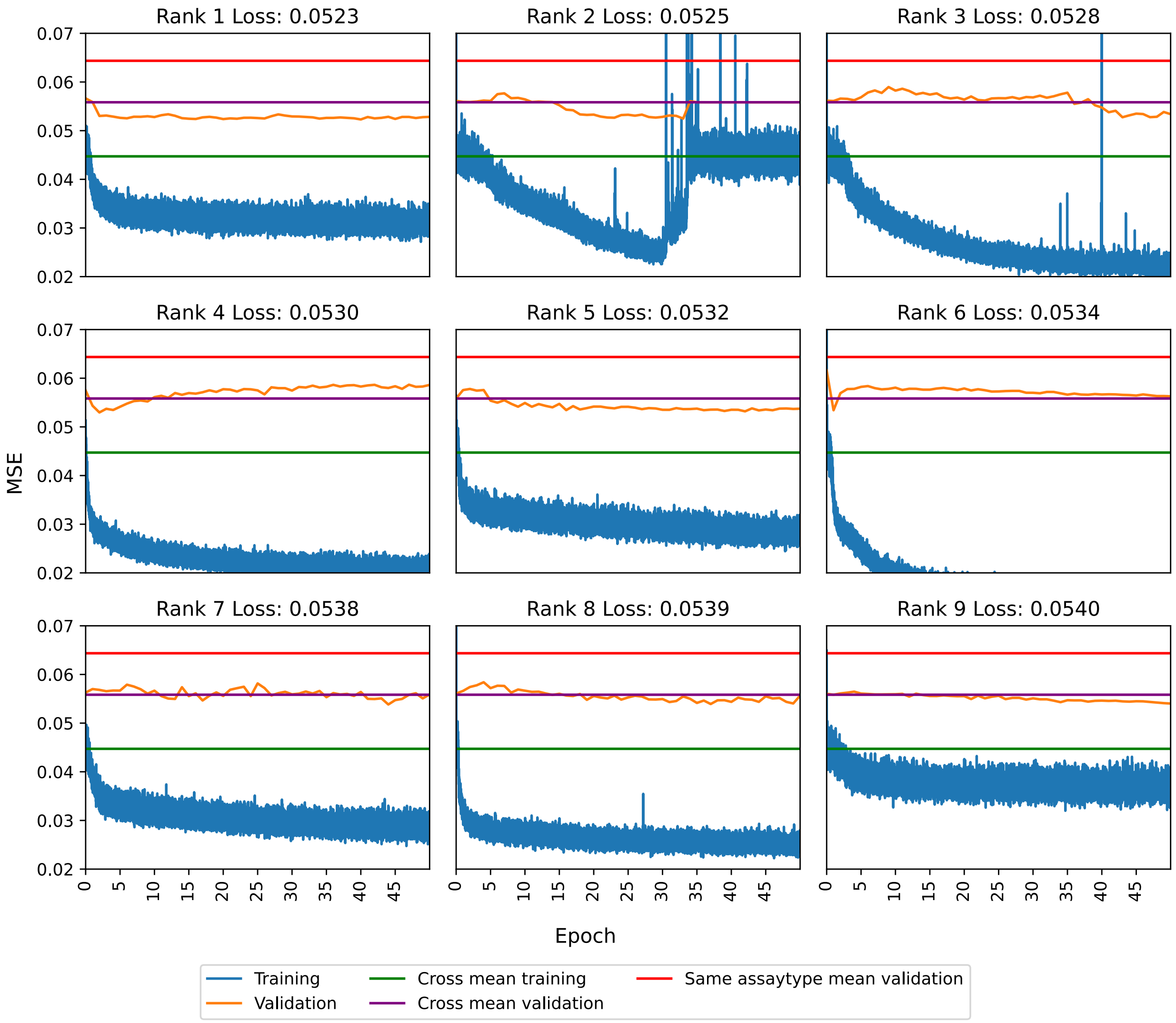


Figure S1: **Loss curves demonstrate sufficient training epochs.** The loss curves for the 9 lowest hyperparameter combinations are shown. The associated hyperparameter combinations are shown in Table S1. The Sphinx training loss (blue), Sphinx validation loss (orange), cross-mean baseline training loss (green), cross-mean validation loss (purple), and same-assay validation loss (red) are shown.
